# Whole Genome Sequence and CAZyme distribution of the cellulase hyper producing filamentous fungus *Penicillium janthinellum* NCIM 1366

**DOI:** 10.1101/2021.06.17.448855

**Authors:** Meera Christopher, Athiraraj Sreeja-Raju, Prajeesh Kooloth-Valappil, Amith Abraham, Digambar Vitthal Gokhale, Rajeev K. Sukumaran

## Abstract

*Penicillium janthinellum* NCIM 1366, capable of secreting cellulases that are highly efficient in the hydrolysis of lignocellulosic biomass, was sequenced to understand its cellulolytic machinery. *De novo* sequencing and assembly revealed a 37.6 Mb genome encoding 11,848 putative proteins, 93% of which had significant *BLAST-P* hits. The majority of the top hits (those with over 60% UniProt identity) belonged to *P. brasilianum*. Carbohydrate active enzymes (CAZymes) and other enzymes involved in lignocellulose degradation were also predicted from this strain and compared with those of the industrial workhorse of cellulase production-*Trichoderma reesei* RUT-C30. The comparison showed that the fungus encodes a far higher number of CAZYmes (422) as compared to *T. reesei* RUT-C30 (244), which gives a plausible explanation for its overall effectiveness in biomass hydrolysis. An analysis of the secreted CAZymes and annotated ligninases identified 216 predicted proteins which may be directly involved in the breakdown of lignocellulose.

## 1. Introduction

The continuous cycling of carbon compounds is one of the fundamental activities in nature. Together with bacteria, fungi are responsible for recycling the bulk of recalcitrant polymers such as lignocellulose [1]. Cellulolytic fungi, i.e., fungi capable of utilizing cellulose-containing material, are widespread in nature and belong to diverse subdivisions like *Ascomycetes, Basidiomycetes*, and *Deuteromycetes*. Amongst them, the most studied aerobic fungal genera are *Chaetomium, Coriolus, Phanerochaete, Poria, Schizophyllum, Serpula, Aspergillus, Fusarium, Geotrichum, Paecilomyces, Penicillium* and *Trichoderma* [2, 3].

The efficient degradation of lignocellulose requires a mixture of different enzymes-cellulases, hemicellulases and ligninases-acting sequentially and in tandem. Cellulases have been traditionally categorized into three different classes by the International Union of Biochemistry (IUB) based on their mode of catalytic action-Endo-1,4-glucanases (EGs) (EC 3.2.1.4) that cleave the cellulose chains internally, Exoglucanases (cellobiohydrolases-CBHs) that include cellodextrinases (EC 3.2.1.74), non-reducing end acting CBHs (EC 3.2.1.91) and reducing end acting CBHs (EC 3.2.1.176) that collectively degrade cellulose from free chain ends; and 1,4-β-glucosidases (EC 3.2.1.21) that hydrolyze soluble cellodextrins and cellobiose into glucose [4]. More recently, carbohydrate-active enzymes (CAZymes), which includes lignocellulases, have been classified into different families based on their structural similarities. These include glycoside hydrolases (GH), glycosyl transferases (GT), carbohydrate esterases (CE), polysaccharide lyases (PL) and enzymes with auxiliary activities (AA) such as the oxidoreduction of phenolic carbohydrates [5].

One of the most efficient and best characterized producers of cellulases is *Trichoderma reesei* (teleomorph: *Hypocrea jecorina*) and its hypersecreting mutants RUT C-30 and CL847 [6, 7]. *T. reesei* secretes large amounts of all three classes of cellulases required for the degradation of cellulose. To date, 9 secretory cellulase encoding genes have been discovered and from *T. reesei-* two CBHs, five EGs, and two BGLs [8–10]. One of the inherent drawbacks of the *T. reesei* cellulase system is its low content of BGLs (~0.5%of the total secreted cellulases), which leads to insufficient degradation of cellobiose and consequently causes feedback inhibition of the endo- and exo-glucanases. To overcome this, commercially available cellulase cocktails, such as those from Novozymes (C-Tec™) and Genencor International Inc. (Accellerase™) rely on the supplementation of heterologous BGLs to improve saccharification efficiency [11, 12].

In the last 20 years, the ascomycete *Penicillium* has emerged as a viable producer of cellulases [13, 14] and sometimes bettering the established industrial work horse for cellulase production *Trichoderma reesei* RUT C30 [15]. Although this genus is traditionally known for industrial drug production, many species, such as *P. pinophilum* [16], *P. brasilianum* [17], *P. citrinum* MTCC 6489 [18], *P. decumbens* [19] and *P. echinulatum* [20, 21], are now known to have extracellular enzyme systems with good lignocellulose hydrolysis potential. Most of these species possess an extensive array of cellulases and hemicellulases that have good stability at 50 □C, a feature that is essential for industrial applications in cellulose hydrolysis. Additionally, in comparison to the enzyme complex of *T. reesei*, some species, such as *P. echinulatum*, have been reported to show more BGL activity in relation to the overall cellulase activity [22].

Previous studies using *P. janthinellum* NCIM 1366 (hereafter referred to as PJ-1366) and its mutants have reported that its cellulases are very efficient in hydrolyzing biomass, including highly recalcitrant lignocelluloses like cotton stalks, at low enzyme loadings [23]. Profiling of the enzyme activities also revealed higher amounts of endoglucanases and ß-glucosidases (BGL) per milliliter of the enzyme as compared to *T. reesei* cellulase. Supplementation of the crude *P. janthinellum* cellulases with BGL from *Aspergillus* species has led to enhancements of its biomass hydrolyzing potential; and in some combinations, it has performed at par or better than the world’s best commercial enzymes for biomass hydrolysis [24]. A recent comparison of secretomes by Sreeja-Raju *et al*. [25] has also revealed that PJ-1366 secretes a higher number of CAZymes on induction, as compared with *T. reesei* RUT-C30 (henceforth given as TR-RC30).

With the implementation of a bio-based economy becoming a logical and necessary option due to environmental and economic concerns, there has been a rising focus on the conversion of cheap and abundantly-available lignocellulose-rich crop residues into a variety of commodity and specialty chemicals and materials, in addition to potential liquid transportation fuels like bioethanol. However, the first step in any of these processes is the biochemical conversion of cellulosic biomass into fermentable sugars by using enzymes, micro-organisms, or other catalysts. Enzymatic transformation of cellulose and hemicellulose into monosaccharides is an effective and sustainable choice, however, worldwide the commercialization of biomass saccharification technologies have proved to be difficult due to the prohibitively high cost of the hydrolases used.

In this scenario, the genome of PJ-1366 has been sequenced, proteins have been annotated using the UniProt Fungi database, and available databases on CAZymes have been used to identify the putative lignocellulose-degrading enzymes of this fungus. This knowledge base will serve as a platform for characterizing the potent cellulase system of this strain, and using it for the development of better biomass hydrolyzing enzyme mixtures.

## 2. Results

### 2.1. Strain confirmation and phylogenetic analysis

*BLAST-N* analysis of the ITS region identified the isolate as *Penicillium janthinellum*. The highest similarity (query coverage-98%, percent identity-99%) was with the ITS region of *Penicillium janthinellum* strain NZD-mf104 (Accession No. **KM278042.1**). The slanted cladogram generated for the phylogenetic analysis showed that PJ-1366 is developed from a basal taxon that also gave rise to *P. brasilianum*, and evolved separately from *T. reesei*. As per the analysis, PJ-1366 is part of a sister taxa to currently reported *P. janthinellum* strains (Fig. 1).

**Fig. 1.**
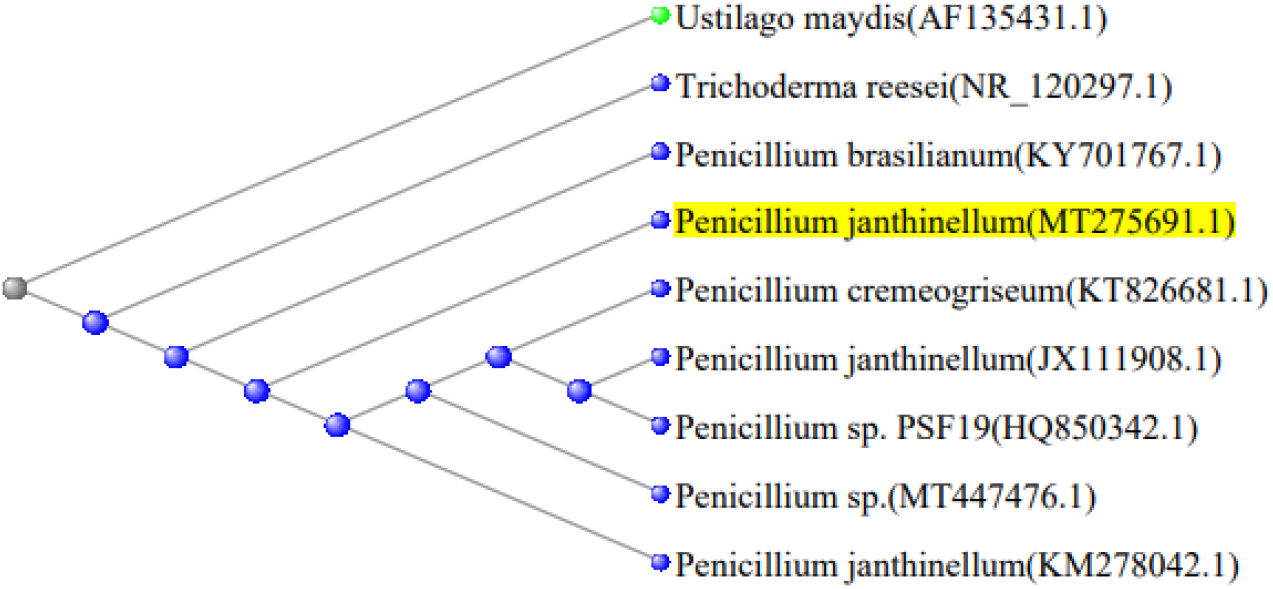
Phylogenetic analysis showing relation of PJ-1366 ITS region with reference sequences from NCBI GenBank. *T. reesei* (ascomycete) and *U. maydis* (basidiomycete) were used as the outgroups

### 2.2. Whole genome sequencing by NGS, de novo assembly and annotation

A total of 30,693,398 raw reads corresponding to 7673.34 megabases, with approximately 100-fold genome coverage were generated by paired-end sequencing. The average base quality was above Q30 (error-probability <= 0.0001) for 93.4% of bases. After data pre-processing, the final data set used for assembly contained 30,055,390 reads covering 7080.7 megabases.

The final assembly consisted of 1773 contigs, with an N50 of 71,572 bp. Based on the assembly, the genome size was estimated to be 37.62 Mb, with a GC content of 50.72%. The genome assembly was deposited in NCBI as a whole genome shotgun sequencing project with accession no. **NPFE00000000.1**

Gene prediction servers predicted 11,848 genes from the genome of PJ-1366. The total length of the coding sequences was 18,352,632 bp, which corresponds to 48.78% of the genome. Out of this, 11,036 predicted proteins (93.14% of total predicted proteins) had significant *BLAST-P* matches with protein sequences in the Fungal UniProt Knowledge Base. Around 94% of the proteins found using *BLAST-P* had a confidence level of at least 1e^−50^, which indicates a high level of protein conservation. Nearly 89% of the predicted proteins had a BLAST similarity score of more than 60% at the protein level with existing proteins in UniProt Fungi. The top *BLAST-P* hit of each protein was studied and it was observed that the majority of the top hits belonged to *Penicillium brasilianum* sp. (Fig. 2)

**Fig. 2.**
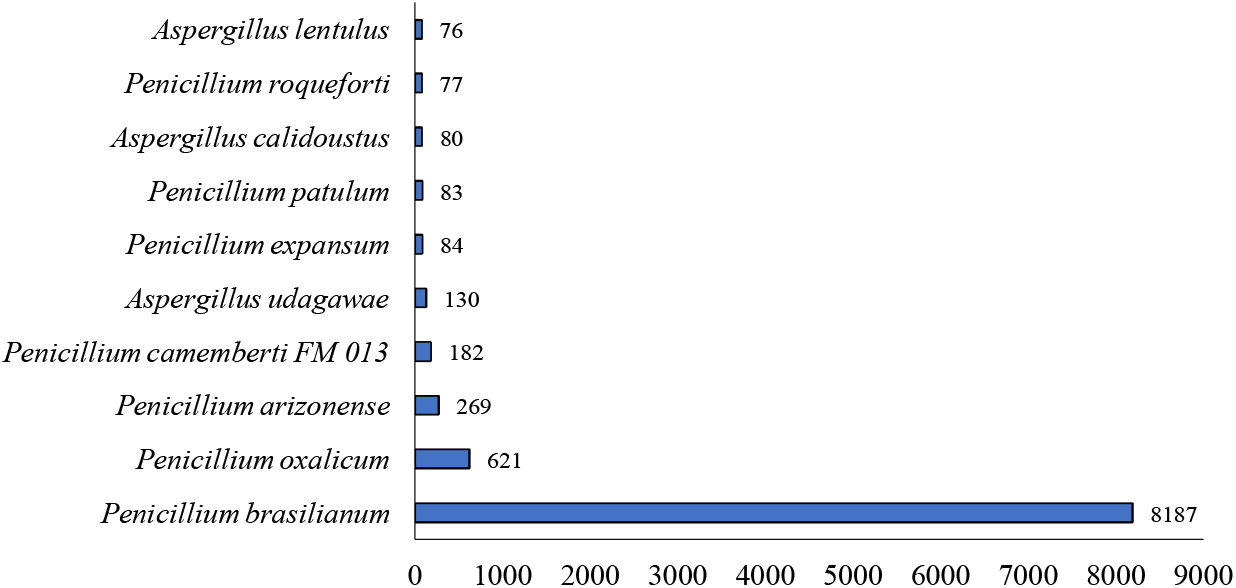
Top 10 organism distribution of proteins of PJ-1366, as obtained by BLAST-P analysis

The gene ontology (GO) terms extracted for the predicted proteins showed that 2721 proteins had predicted molecular functions, 2866 were involved in biological processes and 1992 were cellular components. Of the proteins having molecular function, the majority of them were involved in DNA binding [GO:0003677]. Among the proteins involved in biological processes, 344 were predicted to have a role in transmembrane transport [GO:0055085]. 1364 of the predicted proteins were annotated as being integral component of membrane [GO:0016021] (Fig. 3). Additionally, 206 putative tRNA genes were also extracted by tRNAscan-SE.

**Fig. 3.**
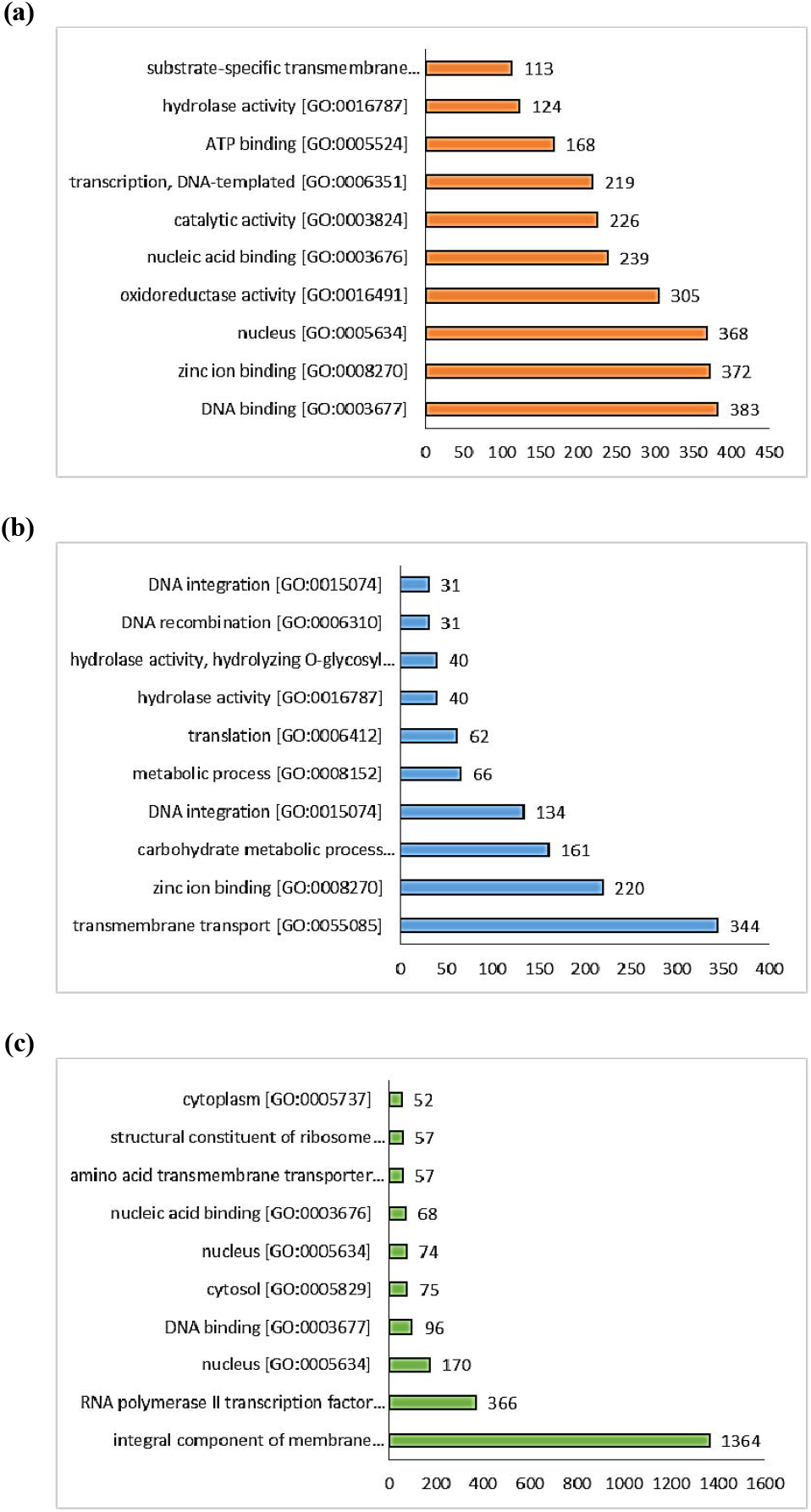
No. of proteins (x-axis) predicted for top 10 GO terms (y-axis) in (a) molecular function, (b) biological process and (c) cellular component

### 2.3. CAZymes of PJ-1366 and TR-RC30

Prediction of CAZymes using the dbCAN server revealed that the genome of PJ-1366 encoded 422 putative CAZymes, as compared to 244 from the genome of TR-RC30. While the number of predicted glycosyl transferases (enzymes that participate in glycosidic bond formation by catalyzing the transfer of sugar moieties from activated donor molecules to specific acceptor molecules) was only slightly higher for PJ-1366, there was a significant difference in the number of enzymes belonging to the other CAZy classes: 1.8x times more glycoside hydrolases (enzymes which hydrolyze the glycosidic bond between two or more carbohydrates or between a carbohydrate and a non-carbohydrate moiety); 12x more polysaccharide lyases (enzymes that cleave uronic acid-containing polysaccharide chains), >2x more carbohydrate esterases (enzymes that catalyze the de-O or de-N-acylation of substituted saccharides) and 1.7x more enzymes with auxiliary activities (Fig. 4).

**Fig. 4.**
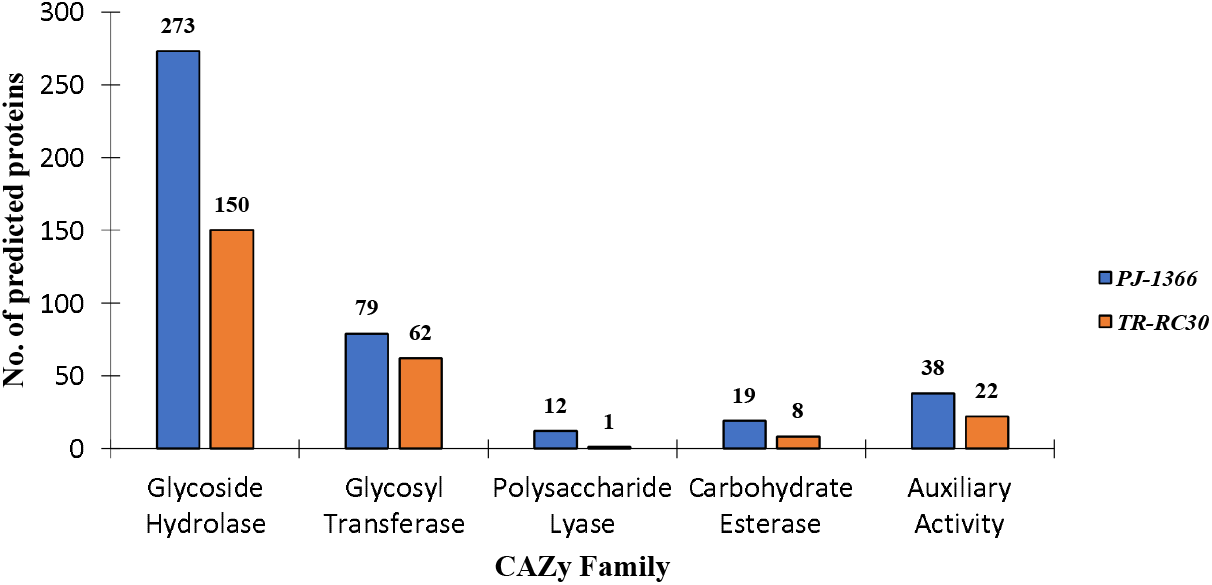
Comparison of putative CAZymes of PJ-1366 and TR-RC30, predicted using dbCAN (No. of tools = 3)

### 2.4. Extracellular biomass hydrolyzing enzymes of PJ-1366

Further, SignalP analysis of PJ-1366 CAZymes showed that 216 of the proteins were extracellular, and could thus be directly involved in biomass hydrolysis. This comprised of 165 GH proteins, 17 proteins belonging to AA class, 10 PLs, 14 CEs and 10 GT proteins. Of the extracellular proteins, 81 of them had predicted CBMs (carbohydrate binding module) that can bind the carbohydrate ligand and direct the catalytic machinery onto its substrate, thus capable of enhancing the catalytic efficiency of the multimodular CAZyme.

Comparison of the putative secreted CAZymes of PJ-1366 with those of TR-RC30 (131 secreted CAZymes) revealed interesting differences-there is an appreciable difference in the distribution of certain CAZy families. PJ-1366 encodes a far higher no. of AA3 (cellobiose dehydrogenase), AA9 (LPMO), CE2 (acetyl xylan esterase), CE8 (pectin methylesterase), GH13 (alpha-amylase), GH19 (chitinase), GH28 (polygalacturonase), GH3 (beta-glucosidase), GH32 (invertase), GH35 (beta-galactosidase), GH43 (beta-xylosidase), GH5 (endo-beta-1,4-glucanase), GT5 (UDP-Glc: glycogen glucosyltransferase), PL1 (pectate lyase) and PL4 (rhamnogalacturonan endolyase) family genes, while TR-RC30 has a higher no. of GH18 (chitinase), GH64 (beta-1,3-glucanase), GH65 (α,α-trehalase), GH67 (alpha-glucuronidase), GH74 (endoglucanase), GT15 (glycolipid 2-alpha-mannosyltransferase), GT17 (ß-1,4-mannosyl-glycoprotein β-1,4-N-acetylglucosaminyltransferase) and PL20 (endo-ß-1,4-glucuronan lyase) proteins (Fig. 5).

**Fig. 5.**
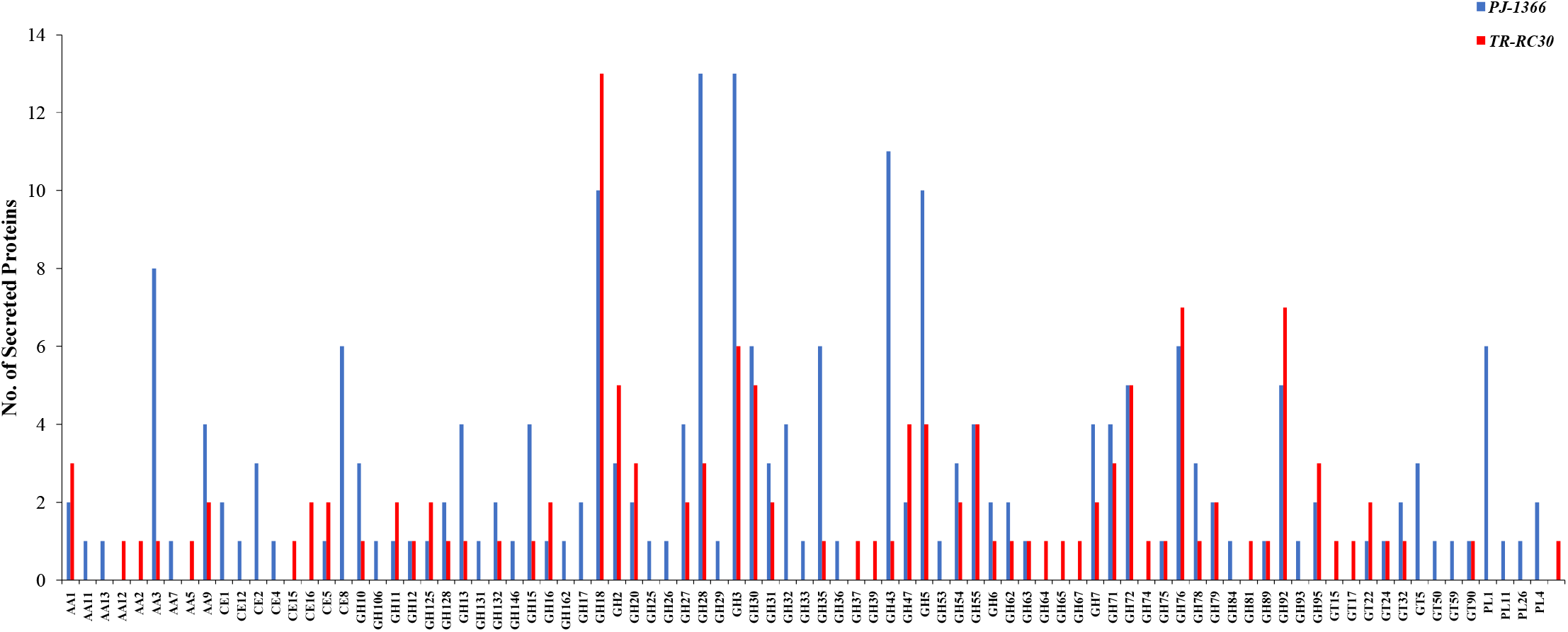
Comparison of extracellular CAZymes of PJ-1366 and TR-RC30

Among the secreted PJ-1366 CAZymes, the most prevalent were GH28 (13 genes), GH3 (13 genes), GH43 (11 genes), GH18 (10 genes), GH5 (10 genes), AA3 (8 genes), and CE8, GH30, GH35, GH76 and PL1 (6 genes each). Six families of auxiliary activity enzymes were represented-AA1 (laccase), AA3, AA7 (glucooligosaccharide oxidase) and 3 families of LPMOs (AA11, AA13 and AA9). Six families of CEs were present, amongst which CE8 was the highest in number. 50 families of GHs were present; the number of GT families represented was 7. As compared to the sole PL of TR-RC30, which is an endo-beta-1,4-glucuronan lyase, PJ-1366 encoded 4 families of PLs-6 genes of PL1, one PL26 (rhamnogalacturonan exolyase) and 3 rhamnogalacturonan endolyases (PL11 and PL4) (Fig. 6). Totally, PJ-1366 encoded CAZymes belonging to 73 unique families, while the CAZymes of TR-RC30 belonged to 59 different families.

**Fig. 6.**
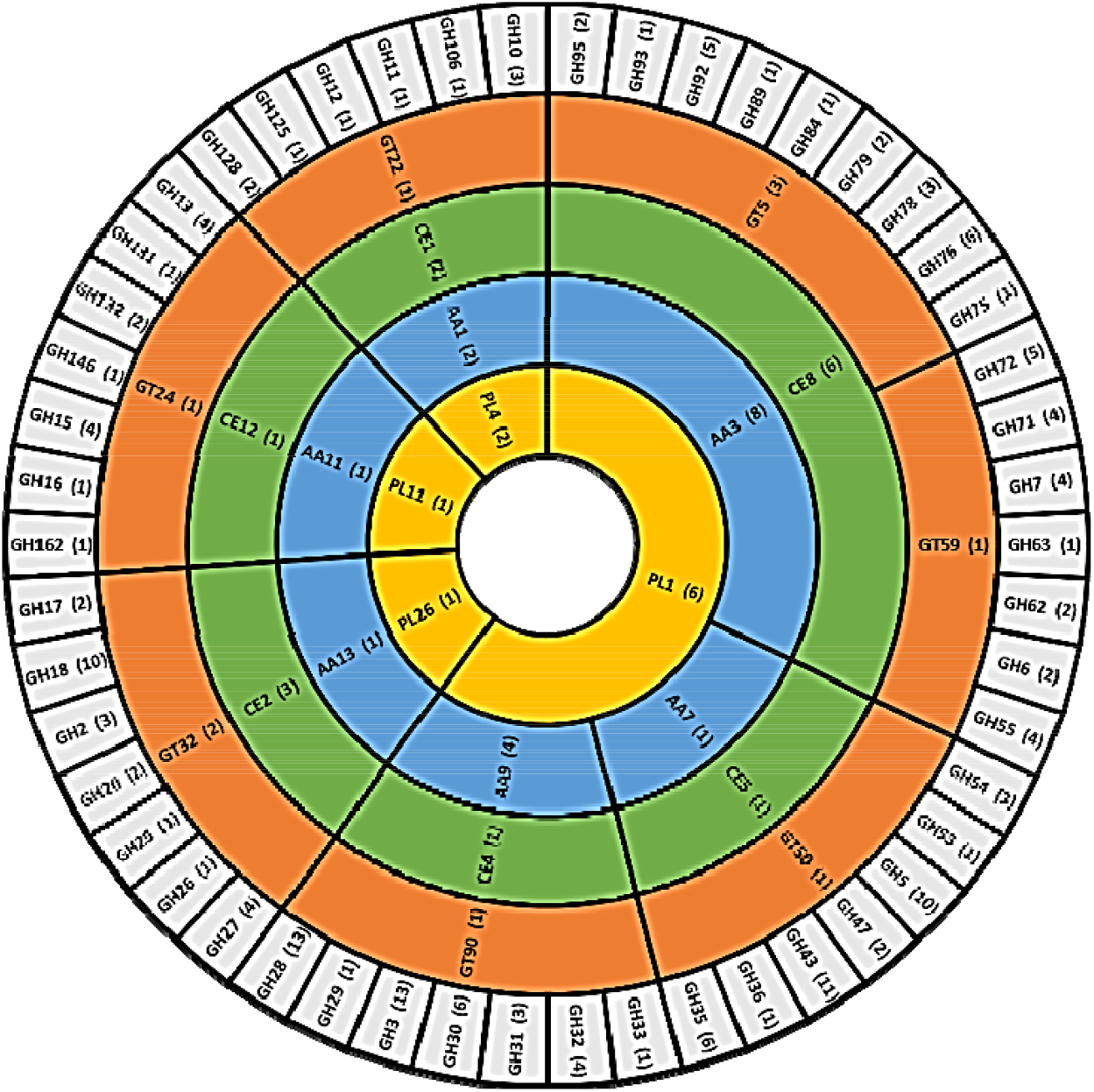
Classification and distribution of extracellular CAZymes of PJ-1366 (Diagram not to scale). Number of putative genes are given within brackets.

From the 165 GHs, 40 sequences were identified which belonged to putative cellulase families GH12, GH131, GH2, GH3, GH30, GH5, GH6 and GH7. *BLAST-P* analysis of these sequences revealed that 19 of them are potential cellulases-4 cellobiohydrolases, 6 endoglucanases and 9 betaglucosidases (Table 1). Out of the remaining 146 sequences, eight sequences were predicted as alpha/glucoamylases, nine sequences were chitinases/chitin synthases, five sequences were GH72 transglycosylases and two sequences were annotated as belonging to the six-hairpin glycosidase superfamily (Supplementary File 1).

**Table 1.**
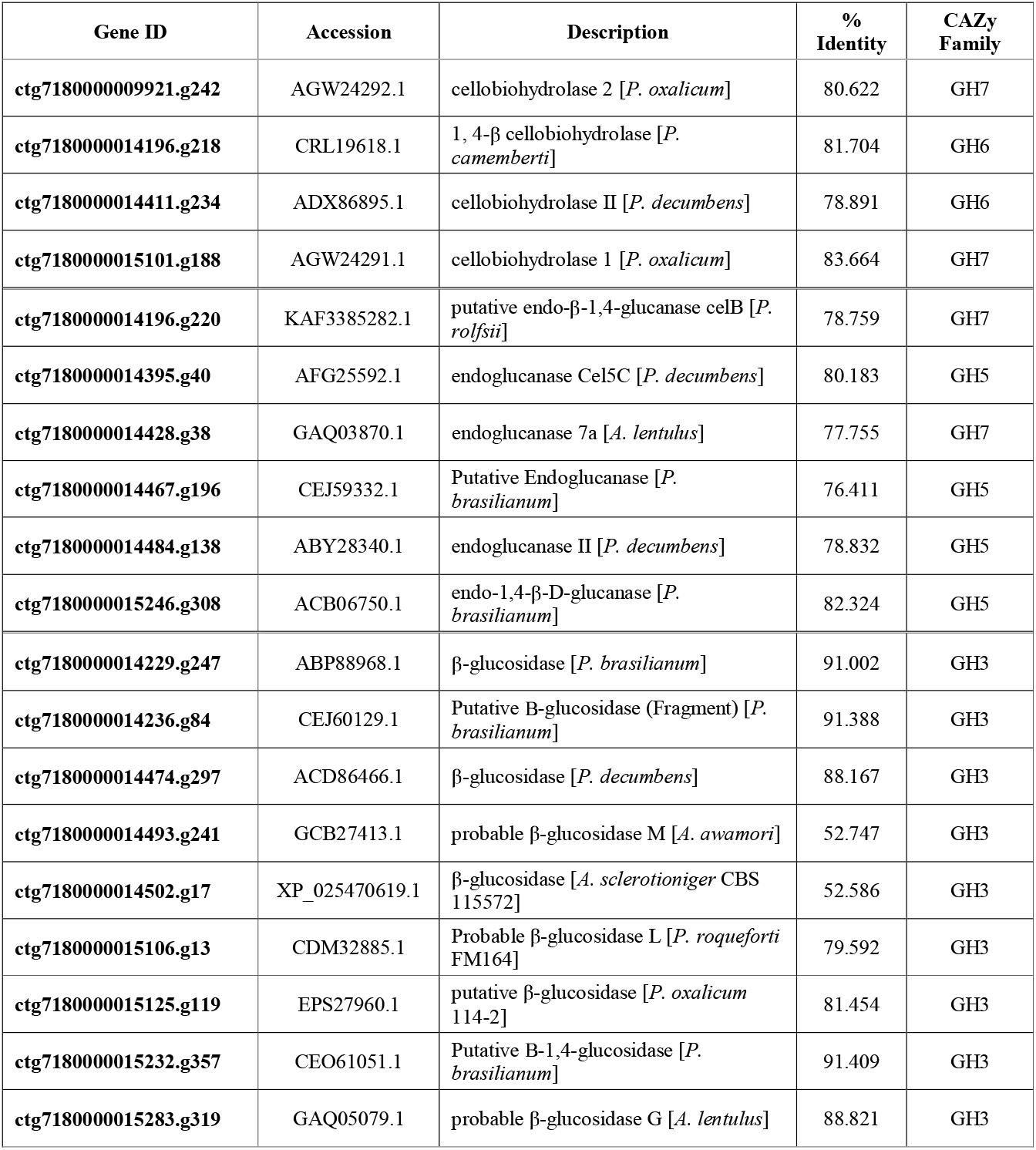
Putative secreted cellulases of PJ-1366

Further, using the available UniProt annotation, 71 sequences were flagged as probable hemicellulases. The most common were xylanases/xylosidases (18 sequences) which cleave the main chain and subsequent oligosaccharides, followed by galactanases/galactosidases (16 sequences). Along with mannosidases (14 sequences), arabinases (9 sequences), rhamnosidases (2 sequences) and galacturonidases (2 sequences), these enzymes are involved in cleaving side chains that bind hemicellulose to cellulose and lignin [26]. Ten sequences encoding polygalacturonases were also present, which, together with pectin lyases (PL1), are involved in the breakdown of pectin [27] (Supplementary File 1)

### 2.5. Ligninolytic enzymes of PJ-1366

Interestingly, the only extracellular ligninolytic enzymes predicted by the dBCAN server were two AA1 laccases (ctg7180000015091.g77 and ctg7180000015373.g58). While laccases (EC 1.10.3.2) facilitate the initial depolymerization of lignin by decarboxylation, demethylation and demethoxylation of its constituent phenolic acid units [28], further degradation requires lignin peroxidases (LiP, EC 1.11.1.14), manganese peroxidases (MnP, EC 1.11.1.13) and versatile peroxidases (VP, EC 1.11.1.16), which are usually grouped in the AA2 family.

There were also no AA2 proteins predicted amongst intracellular CAZymes, though there were a further 2 laccases and one AA5 (oxidase with oxygen as acceptor (EC 1.1.3.-)) protein (Supplementary File 2). An analysis of the annotated proteins of PJ-1366 revealed that there are 5 secretory sequences that are annotated as phenol oxidases and have significant similarity (>60% identity) to fungal laccases in NCBI Protein databases (Table 2). A search for lignin/manganese peroxidases did not give any significant results.

**Table 2.**
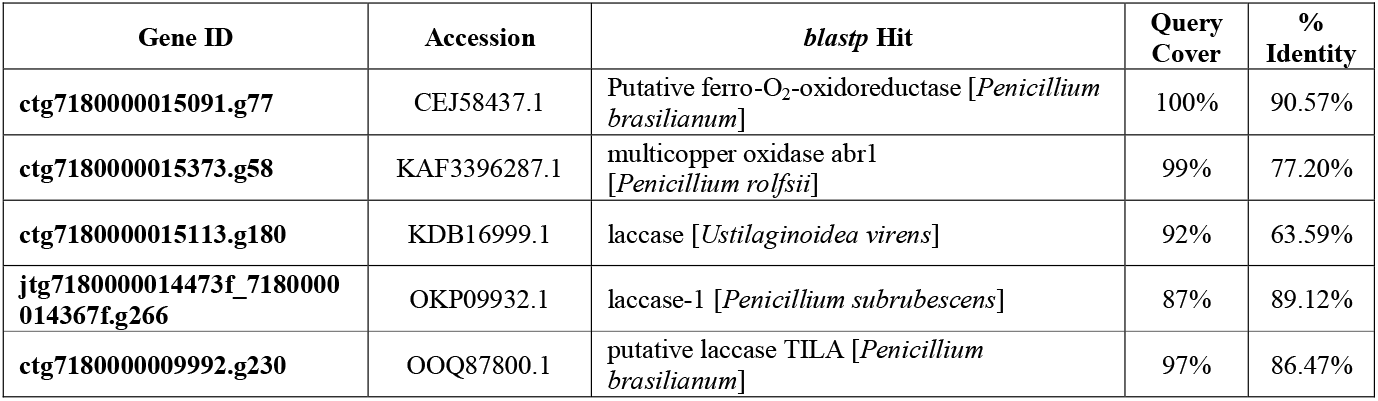
BLAST-P analysis of secretory PJ-1366 proteins annotated as phenol oxidases

## 3. Discussion

In nature, different organisms are capable of degrading lignocellulose; however, the most effective ones are the fungi belonging to *Ascomycota*, like *Aspergillus, Trichoderma* and *Penicillium*. An analysis of the sequenced genomes of some cellulolytic fungi reveals that the size varies from ~ 26 Mb for *P. digitatum* and *P. marneffei* strains [29, 30] to nearly 40 Mb for *T. harzianum* CBS 226.95 [31]. The genome of PJ-1366 is on the larger side, as compared to other sequenced *Penicillium* sp. like *P. chrysogenum* Wisconsin 54-1255 (32.19 Mb) [32] and *P. aurentiogriseum* NRRL 62431 (32.7 Mb) [33]. However, the number of predicted proteins of PJ-1366 (11,848) is comparable to that of *P. aurentiogriseum* (11,476), but higher than that of *P. digitatum* strains (~ 9000). Both the genome size and the number of putative proteins of PJ-1366 is appreciably larger than that of the industrial standard for cellulase production-TR-RC30 (32.7 MB genome encoding 9852 proteins)

Cellulases are multicomponent enzyme mixtures that work in tandem to degrade celluloses and hemicelluloses. In addition to hydrolytic enzymes, an oxidative enzyme system comprised of monooxygenases, dehydrogenases and peroxidases is also involved in boosting the breakdown of lignocellulose [34, 35]. The dbCAN server predicted 422 CAZymes for PJ-1366, a similar number to that reported for *P. decumbens* [19] and *P. verruculosum* [36], but nearly 1.7 times more than the 244 CAZymes reported for TR-RC30

SignalP analysis of the putative CAZymes revealed that more than half of them are extracellular, and thus may be directly involved in the degradation of lignocellulose. Of these, 19 sequences were annotated (using UniProt Fungi and CAZy databases) as cellulases-four CBHs, six EGs and nine BGLs. The CBHs included two each of GH6 and GH7 family proteins. Cellobiohydrolases are considered as the main degraders of crystalline cellulose, with GH6 CBHs being specific towards non-reducing ends of the cellulose chain, and GH7 CBHs attacking the reducing ends [37]. A mutant strain of *P. verruculosum* is known to secrete five different CBHs, but TR-RC30 secretes only two CBHs (one each of GH6 and GH7), which together account for ~80% of its total cellulolytic activity [8, 9]. CBHs are crucial in hydrolyzing pretreated lignocelluloses since the pretreatments carried out to remove the amorphous hemicelluloses and lignin generally result in increased contents of crystalline cellulose [38, 39].

Endoglucanases, on the other hand, cleave the amorphous regions in cellulose chains and are widespread among GH families [40]. Six putative extracellular EGs were identified from PJ-1366, all of which were associated with CBM domains. The presence of a CBM increases substrate binding and generally leads to more thermostable enzymes [41]. Also, the number of predicted EGs in PJ-1366 is higher than both *P. verruculosum* (5) and TR-RC30 (5), which is renowned for its high endoglucanase activity [2]. Even trace levels of EG activity can stimulate CBHs [42], and this might be a reason why, in spite of having similar numbers of EG genes, there is a marked difference in the endoglucanase and FPase activities of cellulases from PJ-1366 and TR-RC30 [25]. Additionally, proteins in GH families are reported to have overlapping specificities, so the function of EGs may also be contributed by the other hydrolases.

PJ-1366 encodes nine putative extracellular GH3 family BGLs, in contrast to the two secreted GH1 and GH3 family BGLs of TR-RC30. GH3 BGLs are known to be functionally diverse, having different pH optima, glucose tolerance abilities, and substrate specificities [43]. Many *Penicillium* species have been reported to have a higher BGL activity than TR-RC30 [44–46], which is advantageous since BGL catalyzes the rate-limiting step in cellulolysis; and therefore the enzyme, if sourced from this fungus may have a bearing on the operational cost of biorefineries [47].

In addition to encoding more cellulases, the number of predicted hemicellulase genes of PJ-1366 (71) also far exceeds the 16 characterized hemicellulases of TR-RC30 [48]. The hydrolysis of hemicellulose is a vital aspect of lignocellulose degradation since hemicellulose typically constitutes 20-40% of cellulose enriched biomass [49]. TR-RC30 has been reported to have a lower secretion of xylan-degrading enzymes, while *Penicillium* species has been reported to secrete a more diversified set of enzymes [6]. The repertoire of PJ-1366 hemicellulases include xylanases and mannanases-that cleave the xylan and mannan main chains of hemicellulose; galaturonases, furanosidases and endo-1,3-glucanases-that digest side chain moieties; and xylosidases, mannosidases and galactosidases-that complete the digestion of oligosaccharides derived from hemicellulose [50]. Together with the BGLs, the GH16/GH72 family proteins (which have probable β-transglycosylation activity), and the ten secreted glycosyltransferases, these enzymes may also be involved in generating soluble inducers for genome-wide induction of cellulases [51, 52]. Studies on bacterial cellulase systems have suggested that while not directly involved in biomass degradation, GT proteins are also invaluable for functionalization (via glycosylation) of the effectors of biomass hydrolysis [53]

Sequences for AA class proteins, which cleave cellulose by oxidative mechanisms, have also been detected, including lytic polysaccharide monooxygenases (LPMOs), which are now understood to create access points for cellulases and hemicellulases [34, 54, 55]. Surprisingly, only some of the enzymes required for lignin degradation were present among the predicted CAZymes of PJ-1366. While a number of lignases have been predicted for TR-RC30 and other fungi, and metal-containing oxidases potentially linked to lignin degradation have been detected in the secretome of TR-RC30 [56], no concrete data exists on their expression patterns. A shortage of information on *Penicillium* peroxidases (550 sequences in NCBI Protein database, as compared to 26881 sequences for *Aspergillus* peroxidases) may also be a contributing factor for the non-availability of structure/sequence data on its lignases. Nevertheless, saprophytic fungi like *Penicillium*, which are involved in recycling lignocellulose residues in nature, must unquestionably be able to efficiently degrade lignins; therefore, there is a need to identify and characterize these proteins by studying the secretomes of lignin-induced fungal cultures.

In conclusion, a cursory analysis of the genome of PJ-1366 and its proteins has revealed that this isolate possesses a complete set of lignocellulose degrading enzymes, which explains its previously reported ability to hydrolyze recalcitrant biomass without the addition of accessory enzymes [23, 25]. The fact that this fungus possesses a more diversified set of lignocellulases than TR-RC30 may be partly due to the fact that *P. janthinellum* is a white rot fungus, unlike *T. reesei* which is generally classified as a soft-rot fungus. By nature of their adaptations to natural habitats, white-rot fungi are capable of degrading all major wood components while soft-rot fungi are only capable of surface degradation of cellulose and hemicellulose. On the whole, the genome assembly and annotation provide a framework for identifying the cellulase machinery of this fungus. In order to fully characterize its lignocellulases so as to use it as a base for strain improvement and development of commercial cellulase cocktails, further studies, including secretome analysis of induced fungal cultures coupled with cDNA profiling, is indispensable.

## 4. Materials and methods

### 4.1. Strain confirmation and phylogenetic analysis

The identity of the strain used for analysis was first reconfirmed by analysis of its ribosomal internal transcribed spacer (ITS) region. 1 x 10^6^ spores were inoculated into potato dextrose broth, and genomic DNA was isolated from the mycelia of 120h old culture by the phenol-chloroform method [57]. Standard protocols were used for the amplification of the ITS region using the conserved eubacterial primers ITS1 (5’ TCCGTAGGTGAACCTGCGG 3’) and ITS4 (5’ TCCTCCGCTTATTGATATGC 3’). The purified PCR product (500bp) was then sequenced and the resulting sequence was compared with the NCBI Nucleotide database using *BLAST-N* [58].

The highest scoring sequences were retrieved from the database as reference sequences and aligned with the ITS sequence of *P. janthinellum*. To assess the relationship between the different strains/species, the taxonomic similarity of isolates was compared using a phylogenetic tree constructed based on the Fast Minimum Evolution method [59], with the maximum sequence difference set at 0.45. The sequences of the ITS regions of *P. brasilianum* (Accession no. **KY701767.1**), *T. reesei* (Accession no. **NR_120297.1**) and the basidiomycete *Ustilago maydis* (Accession no. **AF135431.1**) were used as the outgroups.

### 4.2. Whole genome sequencing by NGS, de novo assembly and annotation

Genomic DNA was extracted as previously described, and treated with RNaseA to remove RNA contamination. The purity and concentration of the DNA were checked using a NanoDrop^™^ ND-1000 UV-Vis spectrophotometer, and the integrity of the DNA was checked by agarose gel electrophoresis.

Next Generation Sequencing (NGS) using the Illumina HiSeq 2500 platform was carried out at AgriGenome Labs Private Limited, Cochin, India. Following paired-end sequencing, the quality of the sequences was checked by assessing various parameters like base quality score distributions, average base content per read and the GC distribution in the reads

All the raw reads were processed before genome assembly. The Illumina adapter sequences were trimmed using Cutadapt v1.8 [60]. All low quality bases (Phred value score <Q30) were eliminated using Sickle v1.33 [61] and duplicate reads were removed using FastUniq [62].

The cleaned reads were subjected to Kmergenie [63] to predict the optimal k-value and assembly size. Contigs were then assembled *de novo* using MaSuRCA v4.0.2 [64]. After the assembly generation, the presence of conserved genes in the assembled contigs was checked using BUSCO v2 [65].

The gene prediction was carried out using an Augustus 3.1 [66] and the predicted genes were annotated using AgriGenome’s in-house pipeline CANoPI (Contig Annotator Pipeline). Briefly it consisted of comparison of the predicted sequences with UniProt Fungi database using BLAST-P program (matches with E-value <= 10^-5^ and similarity score >= 40% were retained for further annotation), organism annotation, gene and protein annotation to the matched genes, gene ontology annotation and pathway annotation.

Finally, tRNA genes were predicted from the contigs using the tRNAscan-SE program [67]

### 4.3. Prediction of CAZymes and identification of lignocellulases

The full repertoire of CAZymes from PJ-1366 were predicted using the web-based dbCAN2 meta server [68]. For comparison purposes, the list of predicted proteins from the genome of *T. reesei (Trichoderma reesei* RUT C-30 v1.0: Project ID-403119) [69] were downloaded from the JGI Genome portal and CAZymes were extracted from it. Only proteins identified as CAZymes using all three tools-HMMER, DIAMOND and Hotpep, were used for further analysis.

The secretory nature of the proteins was predicted using SignalP 5.0 [70]. By comparing the list of secretory CAZymes with the annotated proteins from PJ-1366, a comprehensive list of all lignocellulases, *viz*., cellulases, hemicellulases, lignases and lytic polysaccharide monooxygenases, was prepared.

## 5. Data Availability

The genome sequence is available in NCBI under the BioProject accession number **PRJNA389228**. The contig level assembly is present under accession number **ASM236980v1.** The ITS sequence has been submitted at GenBank with accession no. **MT275691.1**.

## 6. Acknowledgments

MC, ASR and PKV are thankful to the Council of Scientific and Industrial Research (CSIR), Govt. of India for the Junior /Senior research fellowships for Ph.D. studies. This study was partially funded by the Department of Biotechnology, Govt. of India (BT/PR20695/BBE/117/211/2016) and CSIR (MLP 0035: 33/2018/MD-FTT&FTC-ANB). AA is thankful to Kerala State Council for Science, Technology and Environment for a Post-Doctoral Fellowship (001/PDF/2015/KSCSTE). We are thankful to Dr. N. Ramesh Kumar at MPTD, CSIR-NIIST for ITS sequencing service.

